# SAEG: A Novel Deep Learning Architecture for Somatic Alterations

**DOI:** 10.1101/2025.02.21.639424

**Authors:** Luis F. Iglesias-Martinez, Dan Yang Wester, Walter Kolch

**Affiliations:** Systems Biology Ireland, School of Medicine, University College Dublin, Dublin, Ireland; Department of Medical Engineering, Eindhoven University of Technology, Eindhoven, The Netherlands; Conway Institute of Biomolecular & Biomedical Research, University College Dublin, Dublin, Ireland

## Abstract

Somatic alterations, like mutations and copy number changes, driver oncogenesis and cancer progression. Their inhibition has been exploited in the clinic, with several targeted therapies approved for patients with specific mutations or amplifications. However, the response rate of these treatments remains low. The causes are several, ranging from clonal heterogeneity to off target binding. For this reason, CRISPR assays have been developed to study the exact effect of a gene’s deletion. Still, the results from them are puzzling with the same alterations responding different to knockout even in the same cellular context. For this reason, we have developed SAEG, a novel deep learning architecture for somatic alterations in cancer. Our architecture is able to model mutations and copy number alterations and protein-protein interactions to predict if a cell will be susceptible to a gene knockout. SAEG outperforms other models and we show that it learns patterns that can be traced back to the biochemical and biological properties of genes and amino acids.

**Code Availability:** https://github.com/Luisiglm/SAEG

## 1 Introduction

Cancer cells can be reliant on a specific gene for proliferation and survival, this is known as gene dependency (Tsherniak, et al., 2017). This phenomenon can be exploited in the clinic and several inhibitors have been approved to target tumours which harbour somatic alterations that yield gene dependency (Flaherty, et al., 2020; Haslam, et al., 2021; Russo, et al., 2019). However, predicting what somatic alterations make cells dependent on what genes is a non-trivial task.

Broadly, non-silent mutations or amplifications of certain genes are associated with genetic dependency (Haslam, et al., 2021). This is particularly true for a group of genes known as oncogenes, which currently comprise 430 genes known to drive cancer progression (Bernards and Weinberg, 2002; Chakravarty, et al., 2017) Yet, many cells which harbour mutations or amplifications in oncogenes are not dependent on them and consequently do not respond to therapies targeting these genes. In fact, the response rate to targeted therapies in the clinic is as low as 20% (Flaherty, et al., 2020).

Molecular inhibitors often have low specificity and different mechanisms of action that are determined by complex network topologies, including intercalated feedback and feedforward structures (Fey, et al., 2015; Iglesias-Martinez, et al., 2023; Kholodenko, 2015). Thus, drugs can produce hard to interpret results when studying gene dependency. For that reason, CRISPR screens have emerged as a tool to study the effects of gene knockouts in a less confounded fashion (Aguirre, et al., 2016). Still, gene dependency measured through CRISPR screen is not equivalent to or predictive of the effects of a gene mutation or an amplification (De Kegel and Ryan, 2019).

There are several potential causes for cancer cells not to respond to a gene’s inhibition even when it is mutated or amplified. First, mutations can affect a gene in multiple ways depending on what amino acid substitution they produce and on the position which they affect. Some attempts have been made at classifying mutations’ effects using proteins’ conserved regions and structural biology approaches. Yet, connecting structure and function to dependency remains a challenging task. Second, in the case of amplifications, different genes have different dosage effects and a one-fits all model is unrealistic to be a good predictor for gene dependency (Aguirre, et al., 2016; Pugh, et al., 2013) .Finally, other genetic aberrations in addition to the gene of interest can affect a cell’s response to a CRISPR gene knockout or drug mediated inhibition.

Machine learning and artificial intelligence have emerged as potential tools to solve similar problems in other areas (Iglesias-Martinez, et al., 2021; Rukhlenko, et al., 2022). These models can learn complex patterns that can be otherwise hard to elucidate. Large language models have shown high performance in multiple domains (Touvron, et al., 2023; Vaswani, et al., 2017). This includes variant classification, for example the ESM1b model was pre-trained on millions of proteins from the UnifRef dataset and was able to predict germline variant pathogenicity with an accuracy of over 90%(Brandes, et al., 2023) However, this is a large model of 650 million parameters, computing it for several mutations in a cell might be too computational expensive. One of the building blocks of language models are embedding layers that can be used to assign vectors to different word tokens and encode meaning in a mathematical fashion (Vaswani, et al., 2017). Furthermore, sine and cosine embeddings can be used in combination with token embeddings to complement the token vector embedding with its position in the sentence (Touvron, et al., 2023; Vaswani, et al., 2017). Amino acid embeddings have been used both to predict structure and variant class both in evoformer and transformer in AlphaMissense and ESM1b(Brandes, et al., 2023; Cheng, et al., 2023). Yet, these models are computationally expensive and were designed for germline variants instead of somatic.

Here, we propose SAEG a novel architecture, graph neural networks for knockout dependency using similar embeddings to encode somatic alterations and improve gene dependency predictions. Furthermore, we use a graph neural network architecture to also consider somatic alterations in genes interacting directly or indirectly with the gene of interest (Hamilton, et al., 2017; Veličković, et al., 2017). Our results show that our model accurately predicts gene dependency. Interestingly, applying explainable AI techniques showed that the amino acid embeddings learnt by our model reproduce physical properties of the amino acids involved in the mutations without ever being exposed to them explicitly during training.

## 2 Methods

### 2.1 Overview

We designed SAEG, a novel deep learning architecture to handle somatic mutations and copy number alterations, which are the two most common genetic alterations found in cancer. Unlike simpler models, our architecture is able to model point mutations by taking into consideration their position and amino acid substitution. We also include copy number aberrations as they often co-occur with mutations at oncogenes. Finally, our architecture is a graph neural network and thus to predict a cell-line’s dependency on a gene also uses mutations or copy number alterations in genes known to interact directly or indirectly through protein-protein interactions. We used the DepMap CRISPR dataset to train and test our architecture and show that SAEG can accurately predict gene dependency and that it can model the effects of different mutation positions and substitution types.

### 2.2 Graph Neural Network Architecture

We used a graph neural network model to combine copy number aberrations, non-synonymous mutation and protein-protein interaction data to predict gene dependency in cell-lines. Our model is comprised of two different blocks. The first is an embedding block, where gene, mutation position, the original amino acid and the substitution amino acid and copy number are embedded into a learnable vector. The second block is a graph message passing block where genes which are known to interact with each other share information.

### 2.3 Graph Sample Aggregation

We formulate gene dependency as a node regression problem. For each gene in the CRISPR screen, we take its copy number and mutation status and sample a discrete number of genes known to directly and indirectly interact with it using a modified version of the GraphSAGE algorithm (Hamilton, et al., 2017).In the original GraphSAGE algorithm a small number of immediate interactors are selected for each node, and the procedure is repeated for a specific sampling schedule. This works well in large graphs as it reduces the computational expense of evaluating every node in a large network, but in some cases squashes the features of the node of interest. However, somatic mutations and copy number alterations are rare, so sampling each node’s interactors uniformly at random has the risk of yielding nodes with no features. For this reason, for each K nodes being sampled we take every mutated or amplified/deleted gene up to K in the neighbourhood and sample the rest uniformly at random. We followed a [1, 5, 5, 5] sampling schedule yielding neighbourhoods of 126 genes. The p genes sampled, s={g_s1,…,g_sp}, are used to obtain a subgraph that is used in the message passing block described below.

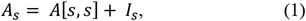

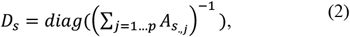

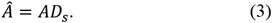

### 2.3 Embedding Block

Each node in our graph neural network passes through a custom embedding layer that maps its features into trainable numeric vectors. As features, we considered somatic mutations and copy number aberrations, and gene (figure 1). We attributed four different features to somatic mutations; a one-hot encoding of whether a mutation is non-silent; and if the mutation is a missense mutation, we used an embedding layer for the original and substitution amino acid and a sine/cosine embedding for the position in the protein where the change occurs (Vaswani, et al., 2017).We used a linear layer on the absolute calls for copy number. All these embeddings are concatenated to build a combined feature vector for somatic alterations. We used 100 features for each embedding, producing 600 features per node.

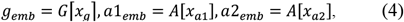

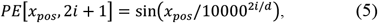

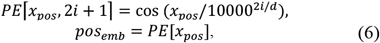

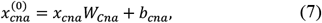

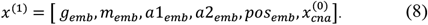

**Figure 1.**
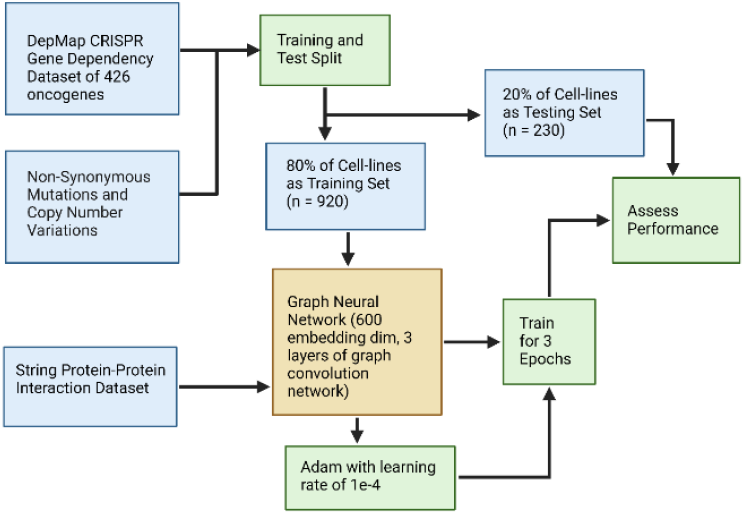
Methods Overview We designed a graph neural network architecture that can use somatic alterations to predict gene dependency, SAEG. We trained our model using the Depmap CRISPR gene dependency dataset focusing on 426 oncogenes. To assess our model’s performance we held out 220 cell-lines that were not used during training.

**Figure 2.**
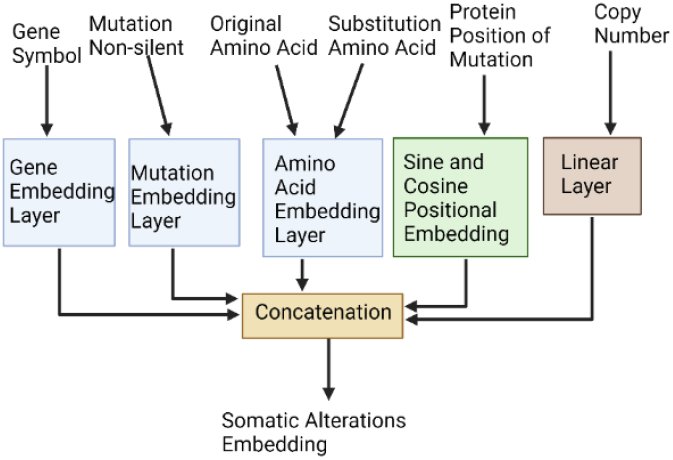
SAEG Somatic Embedding Layer. We built an embedding layer to model somatic mutations and copy number alterations taking into consideration, the position of the substitution, the amino acid substitution and the copy number alterations.

**Figure 3.**
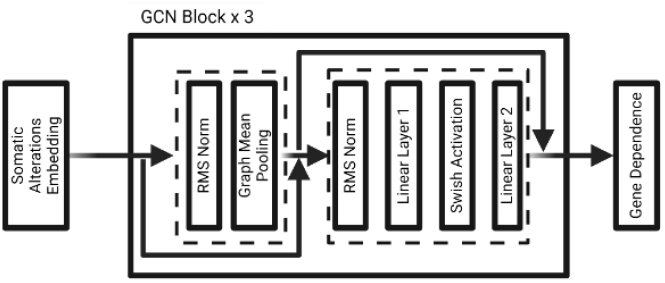
SAEG Architecture. We combined our somatic embedding layer with 3 blocks of graph aggregation layers to predict gene dependency.

### 2.4 Graph Aggregation

We used protein-protein interactions from the STRING database as a graph model (Szklarczyk, et al., 2023). After the somatic mutation and copy number data have been embedded, we use three graph message passing blocks.

At the start of each block, we use graph mean pooling with a residual connection. Then, we use a feed forward neural network with a SiLU activation layer similar to the transformer blocks used in the Llama large language model (Touvron, et al., 2023; Vaswani, et al., 2017). In both cases we use RMS normalization(Zhang and Sennrich, 2019).

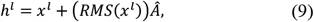

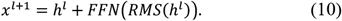

### 2.5 Training

The Depmap CRISPR Gene Dependency dataset was divided into training and testing sets using a 80-20 split (Tsherniak, et al., 2017).We trained our model for 3 epochs using the Adam algorithm with a learning rate of 1e-4 with minibatches of 1 observation and 1 node (Kingma and Ba, 2015). For simplicity, we focused on 426 known oncogenes but used aberration information of any other adjacent genes taken during sampling.

### 2.1 Benchmarking

We compared our model against another graph neural network with one-hot encodings for mutations, and machine learning methods trained on every oncogene without using adjacent genes, namely linear regression and random forests. For the graph neural network, we used 3 layers of mean graph pooling followed by two linear layers, one of the node of interest and one for the graph pooling, concatenated together followed by a PReLU activation layer (Veličković, et al., 2019). For the machine learning methods, we tested both linear and random forest regression using a one-hot encoding for mutations and then random forests using also mutation amino acid substitution and position as variables.

As performance metrics we used correlation because of its interpretability. Since gene dependency has a similar interpretation as a probability measure, we also used accuracy, F1-score and precision assuming that predictions and real values larger than 0.5 are positive labels.

## 3 Results

SAEG had a high correlation (ρ = 0.873, P-value = 0.0) against the test dataset which comprised of 97,980 data points (426 genes in 230 cell lines) (Fig. 4 A). While the linear regression had the highest correlation, and the one-hot encoded graph neural network had the highest F1 score, SAEG outperformed all other methods at accuracy and precision (Table 1). Most methods showed similar performance.

**Table 1.**
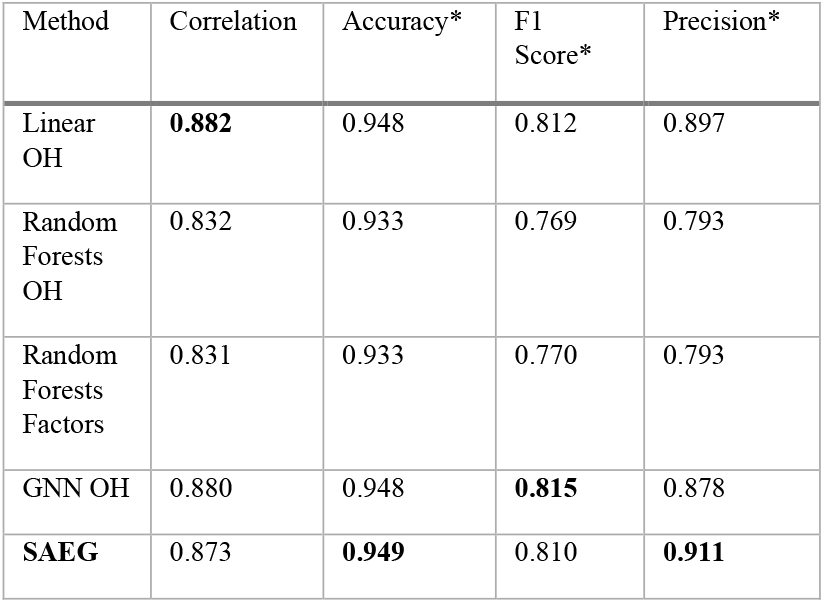
Benchmarking SAEG against other models.

| Method | Correlation | Accuracy* | F1 Score* | Precision* |
| --- | --- | --- | --- | --- |
| Linear OH | **0.882** | 0.948 | 0.812 | 0.897 |
| Random Forests OH | 0.832 | 0.933 | 0.769 | 0.793 |
| Random Forests Factors | 0.831 | 0.933 | 0.770 | 0.793 |
| GNN OH | 0.880 | 0.948 | **0.815** | 0.878 |
| **SAEG** | 0.873 | **0.949** | 0.810 | **0.911** |

**Figure 4.**
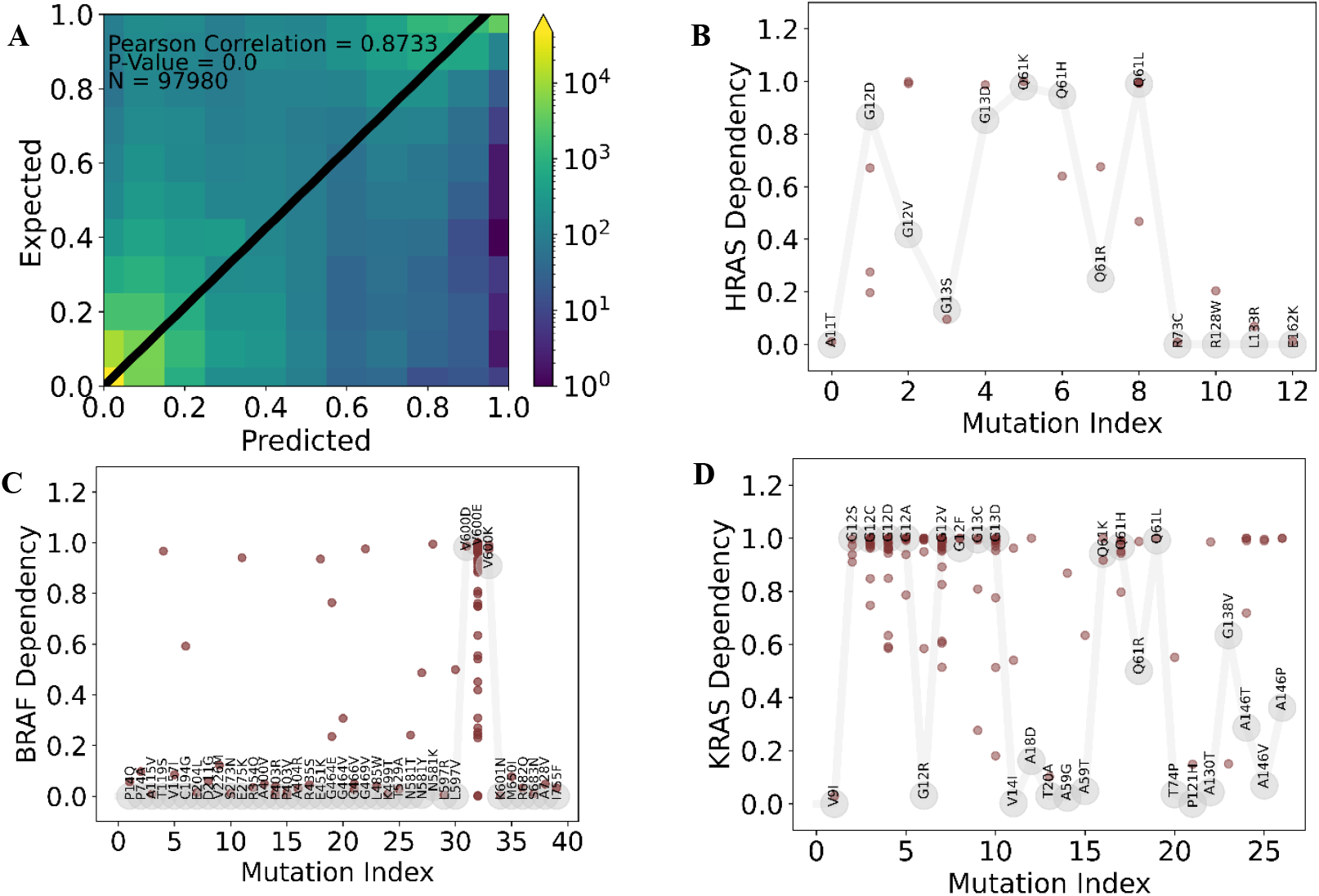
A. Heatmap Predicted Gene Dependency vs Expected Gene Dependecy in the Test Set. SAEG obtained a correlation of 0.87 in the test set indicating a good performance. We used a heatmap because we had all together 97980 points that were hard to represent in a scatter plot. **B HRAS Missense Mutations vs Gene Dependency**. The grey dots represent SAEG’s predictions while the red dots are the observed dependencies across the dataset. In HRAS, only a couple of missense mutations yielded a high dependency score. **C BRAF Missense Mutations vs Gene Dependency**. Most missense mutations did not make the cell’s dependent, while most BRAF V600E mutations did except in some cell-lines. **C KRAS Missense Mutations vs Gene Dependency**. Unlike HRAS, mot KRAS mutations result in dependency as predicted by SAEG and shown in the data. For this experiment, we used mutation data only, and did not use copy number alterations or the mutation status of other genes in our model.

### 3.1 SAEG for Different Point Mutations

To understand why SAEG was predicting dependency in certain cell-line gene pairs, we tested if the somatic alteration embeddings used in SAEG yield different result for each individual mutation. For this experiment, we sampled only KRAS, HRAS and BRAF and did not include the somatic alterations of other genes nor copy number alterations (Fig. 4, B, C and D). Interestingly, in the case of BRAF, SAEG only predicted high dependencies in mutations involving the V600 amino acid which harbours most BRAF mutations, along with most of the data points with high dependency. Most of the rest of the points were classified correctly, except for 7 specific mutations which only appeared one time each in the whole dataset. Similarly, in HRAS and KRAS most of the points associated with high dependency for mutations concerning the G12 amino acid. Unlike in BRAF, our model produced different dependency scores for each. This is consistent with experimental and clinical findings that different RAS mutations have different biological effects. The GTPase activity of KRAS is impaired in mutations in position 12, 13 and 61. The positions in which SAEG gives the highest gene dependency scores. Interestingly, different KRAS mutations also yield different BRAF binding affinities, G12A, V, R and Q61H andL have high BRAf affinity and were assigned higher dependency scores (Huang, et al., 2021).

### 3.2 SAEG for Copy Number Alterations

We also tested if SAEG modelled the copy number alteration differently in each gene. We tested the effects of amplification of two genes, ERBB2 and MYCN, which have different amplification distributions. Interestingly, ERBB2 and MYCN have similar curves (Fig. 5, A and B). However, the predicted dependency grew much faster in ERBB2 compared to MYCN which corresponds to what the data indicate. This is concordant with previous reports indicating a larger dose effect for ERBB2 than MYCN, with MYCN amplified cells harboring significant number of copies of MYCN than ERBB2 amplified cells have of ERBB2(Ghandi, et al., 2019)

**Figure 5.**
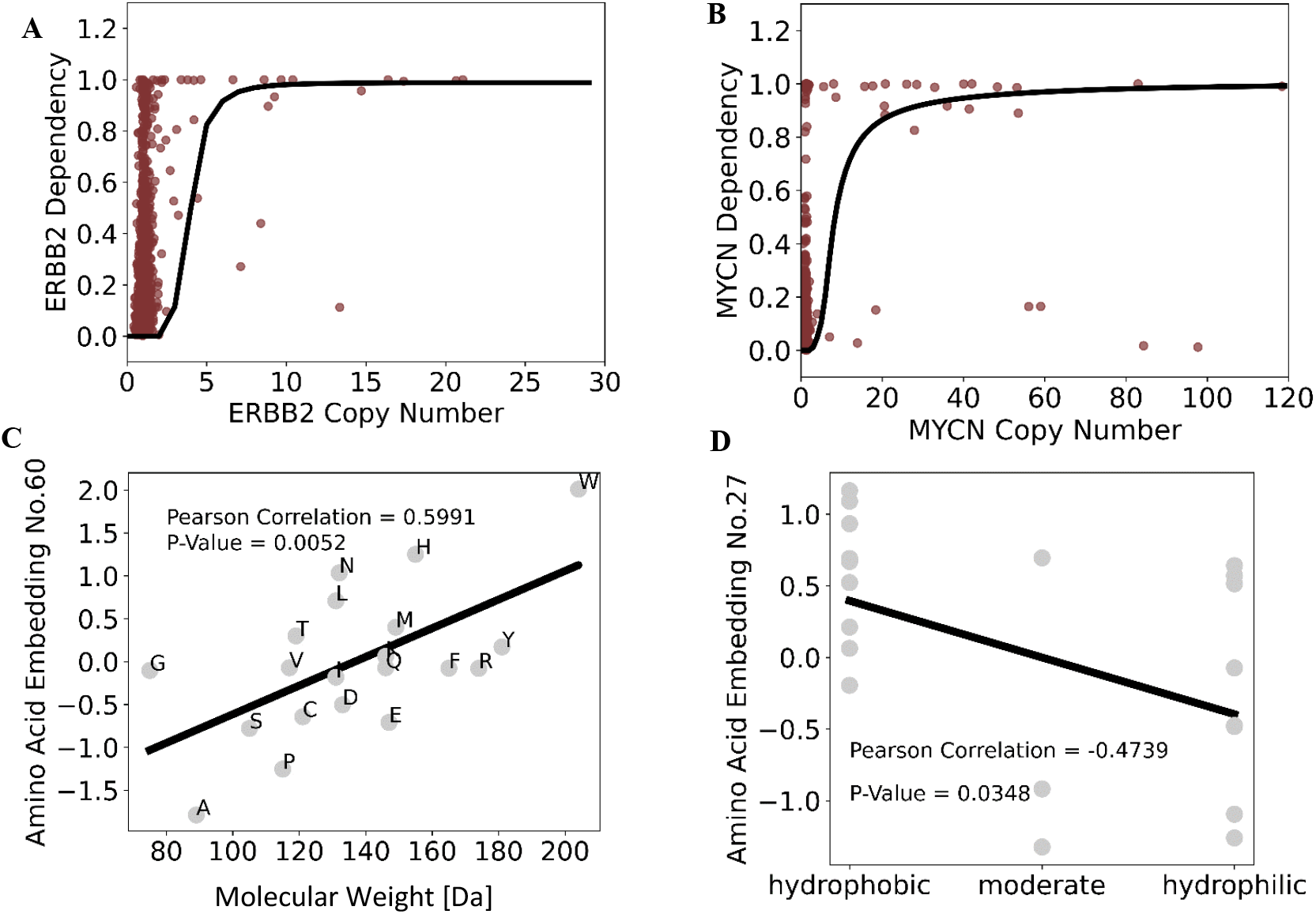
A. ERBB2 Copy Number vs ERBB2 Dependency. The red dots represent data points while the black line are predictions of SAEG given copy number changes in ERBB2 only. **B. MYCN Copy Number vs MYCN Dependency**. The red dots represent data points while the black line are predictions of SAEG given copy number changes in MYCN2. Unlike ERBB2, MYCN was predicted to yield dependency after copy number changes of more than 10, while ERBB2 duplications resulted in high dependency predictions. **B HRAS Missense Mutations vs Gene Dependency**. The grey dots represent SAEG’s predictions while the red dots are the observed dependencies across the dataset. In HRAS, only a couple of missense mutations yielded a high dependency score. **C Amino Acid Embeddings vs Molecular Weight**. Amino acid embeddings 60 was found to be highly predictive of molecular weight (correlation shown in black).. **D Amino Acid Embeddings vs Charge**. Amino acid embeddings 27 was strongly associated with hydrophobicity (correlation lien shown in black).

### 3.3 Modelling Alterations in Interacting Genes

We performed an ablation experiment to see the effect of neighbouring genes (table 2). The results show that the highest performance comes at the largest depth where we sampled genes that were up to 4 edges away from the central node. However, even the model with no neighbours performed well achieving a correlation of 0.855.

**Table 2.**
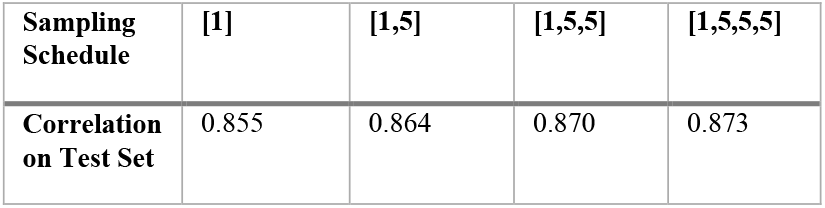
Importance of Neighbourhood Aggregation for SAEG.

| Sampling Schedule | [1] | [1,5] | [1,5,5] | [1,5,5,5] |
| --- | --- | --- | --- | --- |
| Correlation on Test Set | 0.855 | 0.864 | 0.870 | 0.873 |

### 3.4 Amino Acid Embeddings

SAEG uses a somatic alteration embedding layer that takes into consideration the amino acids involved in the substitution. We scanned the learnt embedding vectors of each amino acid to see if it corresponded to their physical and chemical properties. Interestingly, the 60th feature of the embedding vector was strongly associated with molecular weight (ρ = 0.5991, P-value = 0.005) while the 27th feature was correlated with hydrophobicity (ρ = 0.4739, P-value = 0.0348 ).

## 4 Discussion

Our architecture SAEG is able to model accurately gene dependency produced by somatic mutations and amplifications. Our model captures some of the intricacies brought by the differences in mutation positions in different genes. Interestingly, all parameters were shared in our model, meaning that every gene dependency prediction changed by using a gene embedding layer.

Our model was able to reproduce some known differences between mutation positions in BRAF, KRAS and HRAs and in copy number amplifications between ERBB2 and MYCN. Furthermore, our amino acid embeddings were associated with physical and chemical properties that the model was not exposed to, showing that it is capturing some real structural qualities of each mutation. More accurate predictions might be made including protein structure information; however, our model offers a simpler, less computational expensive alternative that can yield precise results(Cheng, et al., 2023). Finally, we did not take cell lineage into account and only modelled other somatic alterations than the gene of interest if they were adjacent in a protein-protein interaction graph. Further improvements could be made by including cellular context and distant somatic alterations.

